# Evolutionary History of *Treponema pallidum* subsp. *pertenue*: Insights into Host Ancestry and Cross-Species Transmission

**DOI:** 10.1101/2025.02.25.640066

**Authors:** Sabrina Karolaine Araújo Sousa de Lima, Tetsu Sakamoto

**Affiliations:** Master’s program in bioinformatics - UFRN; Bioinformatics Multidisciplinary Environment - IMD/UFRN

**Keywords:** Genetic recombination, Yaws, Molecular phylogeny, Genome

## Abstract

*Treponema pallidum*, a bacterium from the Spirochaetota phylum, is responsible for treponematoses, such as syphilis, yaws, bejel, and pinta. Different subspecies of this bacterium cause each of these diseases. This study focuses on *Treponema pallidum* subsp. *pertenue* (TPE), which causes yaws in humans and is primarily transmitted through direct contact with skin lesions, mainly affecting children and preadolescents. If untreated, it can progress to severe deformities in bones and cartilage. During the 20th century, significant progress was made in the eradication and control of this disease. However, in recent decades, there has been an increase in the number of reported cases. Until recently, it was believed that this subspecies affected only humans, but recent studies have identified that non-human primates (NHPs) have also been naturally infected with TPE. Considering the impacts of this disease on both humans and other species, TPE has become a focus of surveillance and scientific investigation. This study aims to clarify the relationship between infection in humans and other primate species, contributing to a better understanding of transmission dynamics and potential control and prevention strategies. To achieve this, we used genome sequences from 58 TPE strains (24 from humans and 19 from NHPs) available in public repositories and applied phylogenetic analyses and computational tools for recombinant region detection. The phylonetic analysis demonstrated a rapid expansion of the subspecies at the base of the tree and clustered the samples into nine groups. No groups contained both human and NHP samples, indicating that interactions between the two sample groups are rare or nonexistent. Recombination analyses detected only one region where an NHP sample may have recombined with a human sample. The application of the molecular clock in phylogenetic inference indicated a recent origin of TPE in 1885. The study of trait evolution further suggests that the host of the most recent common ancestor of TPE was humans. The analyses conducted in this study provided deeper insights into the spread of the disease and its interaction across different species.

## 1. Introduction

*Treponema pallidum* (phylum *Spirochaetes*, order *Spirochetales*, family *Spirochaetaceae*) is a bacterial species responsible for causing treponematoses, which are progressive pathogenic infections that include syphilis, yaws, bejel, and pinta. Each subspecies of this pathogen is directly associated with different diseases. The subspecies *endemicum* (TEN) causes endemic syphilis (bejel); *T. pallidum* subsp. *pallidum* (TPA) causes venereal syphilis; *T. pallidum* subsp. *pertenue* (TPE) causes yaws; and *T. pallidum* subsp. *carateum* causes pinta (MITJA; SMAJS; BASSAT, 2013). Among these four highly morbid diseases, venereal syphilis stands out because it has a high prevalence globally and is transmitted through sexual contact, while the other three are tropical neglected diseases which are spread through close personal contact. However, yaws disease caused by TPE has been highlighted in recent studies because it is the only subspecies found to infect other species of non-human primates (NHPs), making interspecies transmission of this disease possible (Knauf et al, 2018).

Yaws is more common in children from low-income rural communities in tropical areas of Africa, Asia, Latin America, and the Pacific. Although easily curable in its early stages, advanced cases can cause irreversible bone deformities. In 1952, the World Health Organization (WHO) launched the first global campaign to eradicate yaws, distributing penicillin-based antibiotic injections. This led to significant progress in combating the disease: 50 million people received treatment, reducing the global disease burden by 95%. WHO had projected yaws eradication by 2020, but this goal was not achieved (PAHO, 2018). Between 2008 and 2015, nearly 462,000 cases of yaws were reported to WHO in 12 endemic countries located in Africa, Southeast Asia, and some areas of the Pacific. Children under the age of 15 account for more than 75% of reported cases (PAHO, 2018). In 2020, more than 300 cases were confirmed, and over 80,000 people with suspected clinical cases of yaws were recorded in 15 countries, but this is likely an undercount, since many countries stopped reporting cases to the WHO (PAHO, 2023). There are at least 76 previously endemic countries where the disease status remains unknown (JOHN et al., 2022).

The current perspective is that the disease will be eradicated by 2030. However, new obstacles have been brougth to light by recent researches that may make the eradication programme challenging (BOLLINGER, R., 2024). One of these is the emergence of antibiotic resistant strains and the other, which is the focus of this work, is the existence of a TPE reservoir in some NHP species. For nearly a century, humans were considered to be the exclusive hosts of the bacterium that causes yaws. In fact, several wild population of African NHPs were described to exhibit skin ulcerations suggestive of *Treponema* infections, and antibodies against *T. pallidum* (LEVRÉRO et al., 2007, KNAUF et al., 2012, KNAUF et al., 2018). However, it was only after the genome sequencing that it was discovered that the strains infecting NHPs are TPE, suggesting the existence of a natural reservoir for the infection.

The recognition of *T. pallidum* infections in non-human primates (NHPs) has led to the hypothesis that human treponematoses may have a zoonotic origin (KNAUF; LIU; HARPER, 2013). It is still debated whether human *T. pallidum* results from a single transmission event, continuous transmission, or if these *Treponema* species evolved in parallel with their primate hosts (HARPER et al., 2008). It has been suggested that efforts to eradicate yaws may be hindered by the potential for continuous transmission from a reservoir of these pathogens in NHPs, although there is currently no evidence confirming transmission events between NHPs and humans in the wild (KNAUF; LIU; HARPER, 2013).

These observations suggest that the impact of treponematoses may be broader than initially thought, affecting not only humans but also other primate species (HARPER et al., 2012). The available evidence is insufficient to conclude whether other species are also being infected, highlighting the need for further studies and investigations to fill this gap in scientific knowledge (Knauf et al., 2012). This study aims to deepen the understanding of the dynamics of these diseases in humans and NHPs. We aimed to examine the role of NHP as possible natural reservoirs of the bacterium, as well as the associated zoonotic risks, providing scientific insights for public health policies, biodiversity conservation, and yaws control strategies. To achieve this, we used genomic data of TPE publicly available and applied Bioinformatics tools to identify genetic recombination events between human and NHP samples, and to investigate the origin and the potential ancestral host of the pathogen.

## 2. Methods

### 2.1. Sequencing Data and Pre-Processing

The *TPE* sequencing data used in this study were based on those utilized in the study by JANEČKOVÁ et al., 2023. A total of 80 *TPE* sequencing samples were downloaded from the National Center for Biotechnology Information (NCBI) repository. Raw read data were obtained from 58 samples using the Sratoolkit tool (https://github.com/ncbi/sra-tools), while nucleotide sequences from assembled genomes were obtained from 22 samples via the NCBI webpage. Additionally, nucleotide sequences from assembled genomes of two *TPA* strains (*Nichols* and *SS14*) were also retrieved to serve as an outgroup in the phylogenetic analyses. The sequencing data, both raw read data and assembled genomes, were processed using the Snippy program (https://github.com/tseemann/snippy) which mapped and aligned all sample sequences against a reference genome. Snippy is a tool that aids in the identification of genomic variants, with an emphasis on detecting single nucleotide polymorphisms (*SNPs*) and small insertions and deletions (*indels*). The reference genome used was from the *TPE* strain SamoaD. The quality of the sequencing data was assessed by examining the results obtained from Snippy. Samples that were not mapped to more than 10% of the reference genome were excluded from subsequent analyses. In the end, 43 samples were selected (41 *TPE* samples and 2 *TPA* samples, as listed in Table 1), with 24 samples from *Homo sapiens*, 2 from *Cercocebus atys*, 3 from *Chlorocebus pygerythrus*, 5 from *Chlorocebus sabaeus*, 7 from *Papio anubis*, and 2 from *Papio papio*. The geographic locations of the human samples include Ghana, Solomon Islands, Liberia, Congo, Indonesia, the United States of America, and Samoa. The non-human primate samples originate from Ivory Coast, Tanzania, Senegal, The Gambia, and Guinea.

**Table 1.**
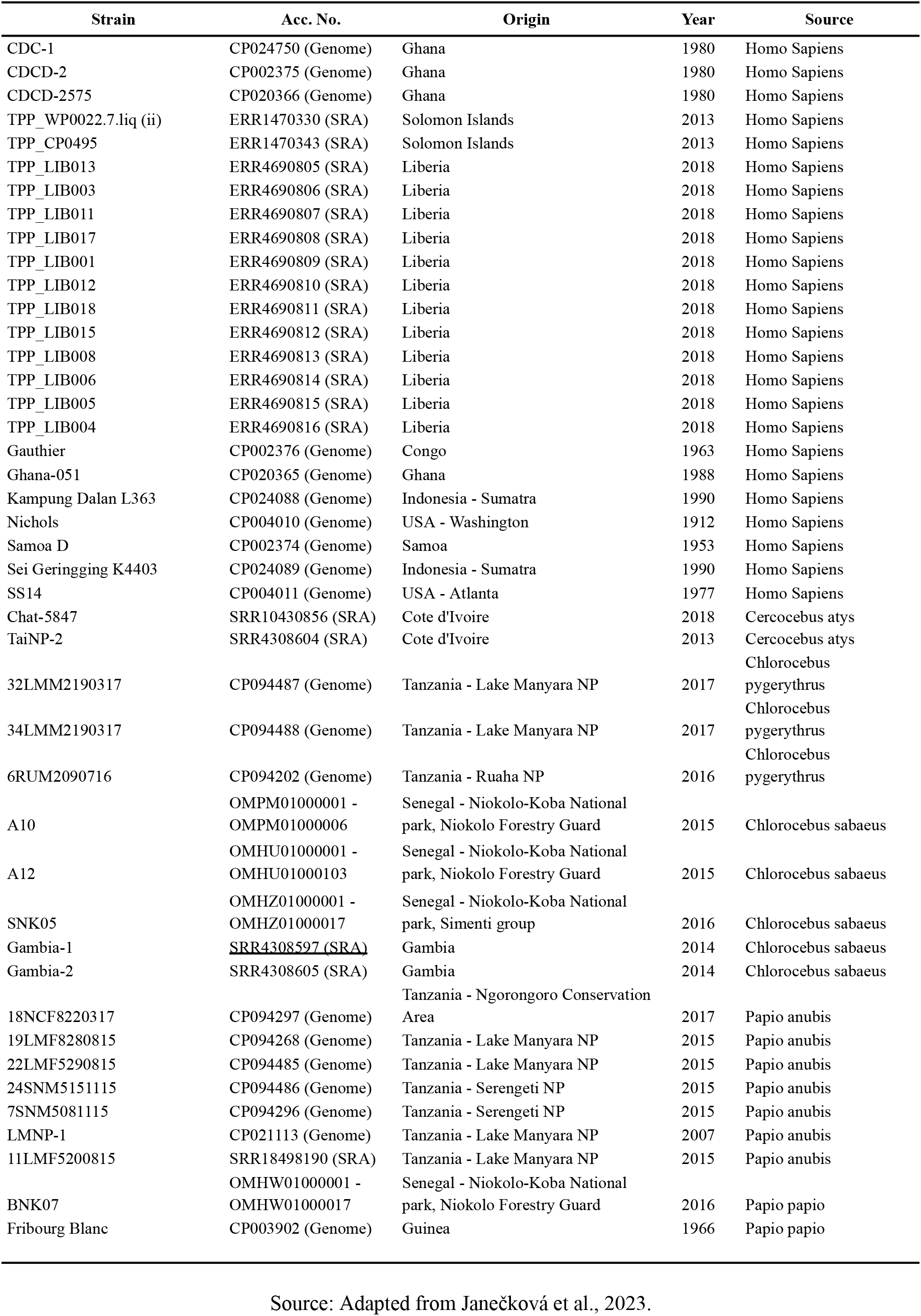
*Treponema pallidum* Samples Used in This Study and Their Metadata.

### 2.2. Phylogenetic Analysis

IQ-TREE (MIN et al., 2020) was used to perform maximum likelihood (ML) tree inference. The sequences used for this analysis correspond to the full genome alignment generated by the Snippy program. The best nucleotide substitution model was selected based on the Bayesian Information Criterion (BIC) using ModelFinder (KALYAANAMOORTHY et al., 2017), which is integrated into IQ-TREE. Statistical support for each branch was calculated using ultrafast bootstrap with 1,000 replications. The final tree was rooted using *TPA* samples and visualized using FigTree (http://tree.bio.ed.ac.uk/software/figtree/).

The temporal and evolutionary analysis was conducted using BEAST, which employs Bayesian inference methods to estimate time-calibrated trees and evolutionary parameters. The input for this analysis included SNP alignment data, the year of isolation, and the host species (*Human* or *Non-Human Primate*) for each sample. The bModelTest package (BOUCKAERT; DRUMMOND, 2017) was used to estimate the most appropriate evolutionary model, while the Optimized Relaxed Clock model (DOUGLAS; ZHANG; BOUCKAERT, 2021) was used to estimate divergence times among taxa. The Coalescent Bayesian Skyline model was applied as the tree prior. Markov Chain Monte Carlo (MCMC) chains were set to run for 200 million iterations, with sampling every 10,000 iterations. Before running BEAST, the Extensible Markup Language (XML) file was modified using beast2_constsites (https://github.com/andersgs/beast2_constsites) to account for the constant sites in the analysis. The choice of the molecular clock model was based on the R^2^ statistic obtained from the root-to-tip regression analysis performed with TempEst (RAMBAUT et al., 2016). For this analysis, the input data included the ML tree generated by IQ-TREE and the isolation year of each sample. The BEAST program was executed twice independently. The Tracer software (RAMBAUT et al., 2018) was used to check for MCMC chain convergence and to evaluate Effective Sample Size (ESS) values for the inferred parameters. The trees generated from the two independent runs were combined using LogCombiner, a tool within BEAST, discarding the first 10% of trees from each chain. Finally, the Maximum Clade Credibility (MCC) consensus tree was generated using TreeAnnotator, also from BEAST, with node heights rescaled based on the mean height of the common ancestor among the sampled trees. The final tree was visualized using FigTree (http://tree.bio.ed.ac.uk/software/figtree/).

The construction of the unrooted phylogenetic network was performed using SplitsTree (HUSON; BRYANT, 2024). In this analysis, only SNP alignment data were used as input. The network was constructed using the Neighbor-Net method, and distances were calculated using P-distance. This tool was employed to identify recombination signals among the samples or phylogenetic conflicts.

### 2.3. Identification of Recombination Signals

Inferences regarding genomic regions that may have undergone recombination among the samples were performed using the Gubbins (NICHOLAS et al., 2015) and FastGear (MOSTOWY et al., 2017) programs. For both programs, the full genome alignment generated by the Snippy program was used as input. For the execution of Gubbins, the input included the full genome alignment of *TPE* samples along with the external samples (*TPA*), as generated by Snippy, as well as the maximum likelihood tree generated by IQ-TREE, rooted using the outgroup samples. Additionally, the program was configured to run with default values, except for the recombination model, which was set to GTRGAMMAIX. For the execution of FastGear, the same full genome alignment used in Gubbins was also provided as input, but without the *TPA* samples. All other parameters were kept at their default values.

## 3. Results

The results of this study show that the genomes of *Treponema pallidum* subspecies *pertenue* (TPE) exhibit low genetic variability. Using the reference genome SamoaD, which is approximately 1.13 Mbp in size, the Snippy tool identified 927 core SNPs, which are variants found at sites present in all samples. The distribution of SNPs along the genome was not uniform (Figure 1), as evidenced by the KS statistic (0.077; p-value: 0.000028), indicating the presence of regions with a higher accumulation of variants, which suggests potential mutation hotspots or regions under greater selective pressure. Among the 927 identified SNPs, two were multiallelic, while the rest were biallelic. The number of transitions and transversions was 732 and 197, respectively. The frequency of each type of substitution is summarized in Figure 2.

**Figure 1.**
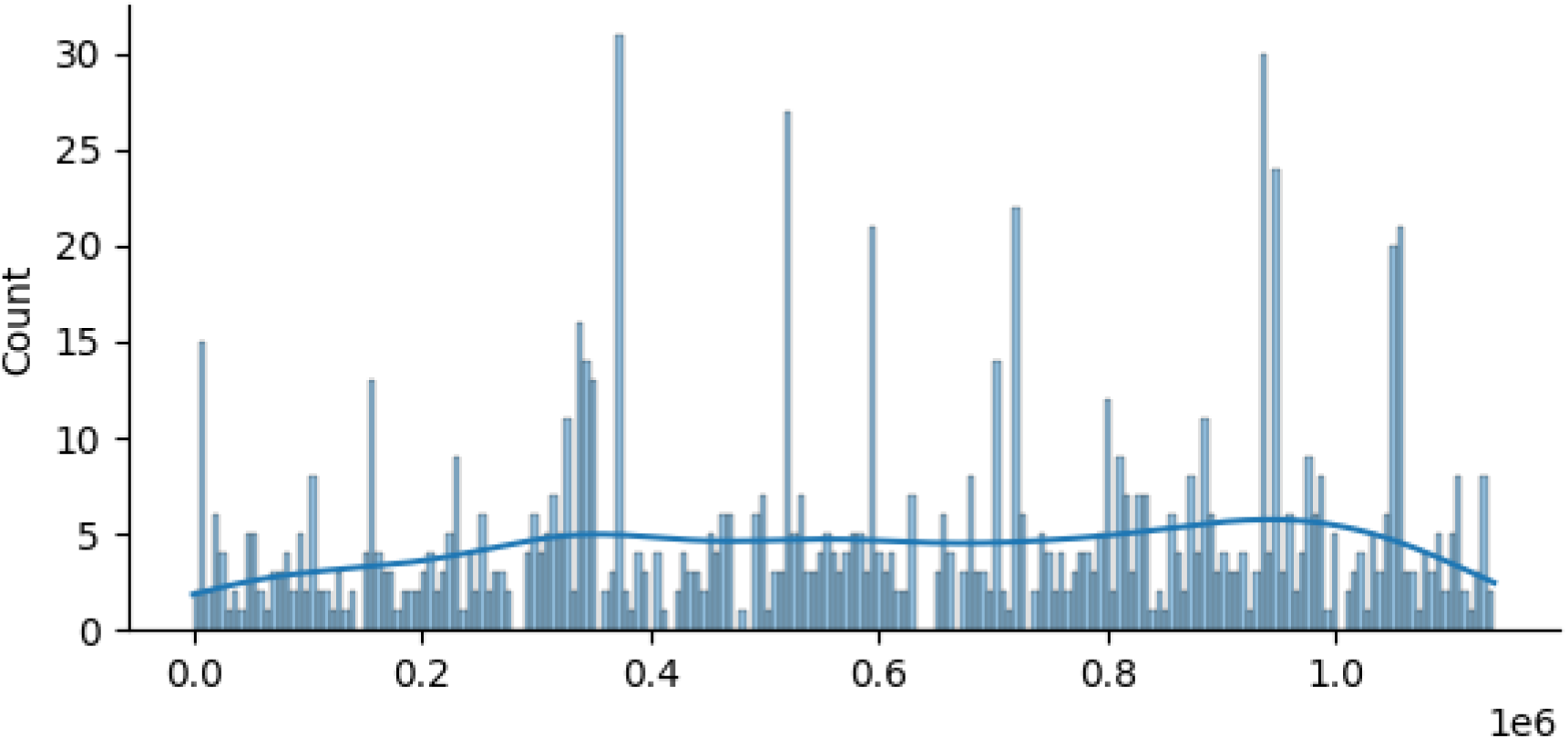
Distribution of SNPs along the *TPE* genome. The X-axis represents the genome positions of the reference sample (*SamoaD*). The Y-axis represents the number of SNPs found within a given range of genome positions. The line represents the estimated probability density function of the SNP distribution, generated using Kernel Density Estimation (KDE).

**Figure 2.**
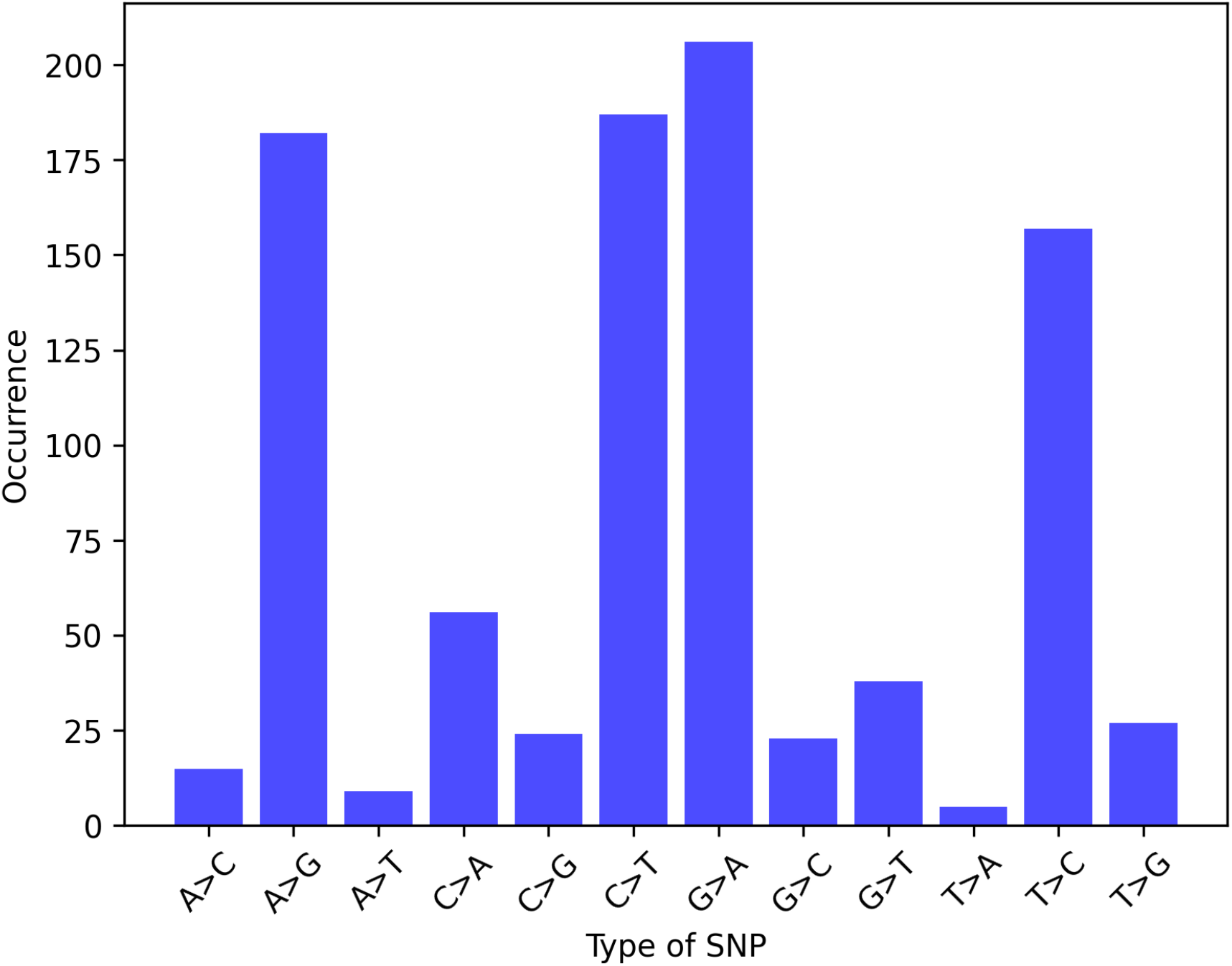
Frequency of Genomic Variations. The X-axis represents the different types of variations present in the genome, while the Y-axis indicates the number of occurrences of each variation in the analyses performed.

### 3.1. Much of the evolution of *Treponema pallidum* subsp. *pertenue* remains unresolved

The phylogenetic analysis performed using IQ-TREE revealed the presence of ten groups, seven of which are monophyletic clades with high statistical support (bootstrap value > 90), while the remaining three consist of a single sample each that did not cluster with any other clades. The tree indicates distinct clustering patterns within the *Treponema pallidum* subsp. *pertenue* species (Figure 2). Among these groups, three were composed exclusively of samples isolated from non-human primates, while the other seven consisted only of samples isolated from humans, suggesting potential transmission barriers between hosts. However, the internal nodes located at the base of the phylogenetic tree exhibited low support values and short lengths, indicating low resolution in the deeper evolutionary relationships within the subspecies.

This result can be explained by a rapid expansion process of the lineage in different hosts or by the occurrence of genetic recombination events between clades, making it difficult to clearly define phylogenetic relationships. These findings highlight the need for further investigations using complementary approaches, such as recombination analyses and evolutionary modeling, to better elucidate the evolutionary history of this subspecies.

### 3.2. Limited Evidence of Recombination Signals Between Samples Isolated from Human and NHP Hosts

To identify potential gene recombination events among *TPE* groups, we first used SplitsTree to infer reticulated trees. The application of network-based methods, such as NeighborNet, allows for the visualization of non-hierarchical clustering, which may indicate complex evolutionary events such as recombination or horizontal gene flow. The generated reticulated tree showed few reticulations among the samples, indicating the absence of recombination events between *TPE* clades (Figure 3).

**Figure 3.**
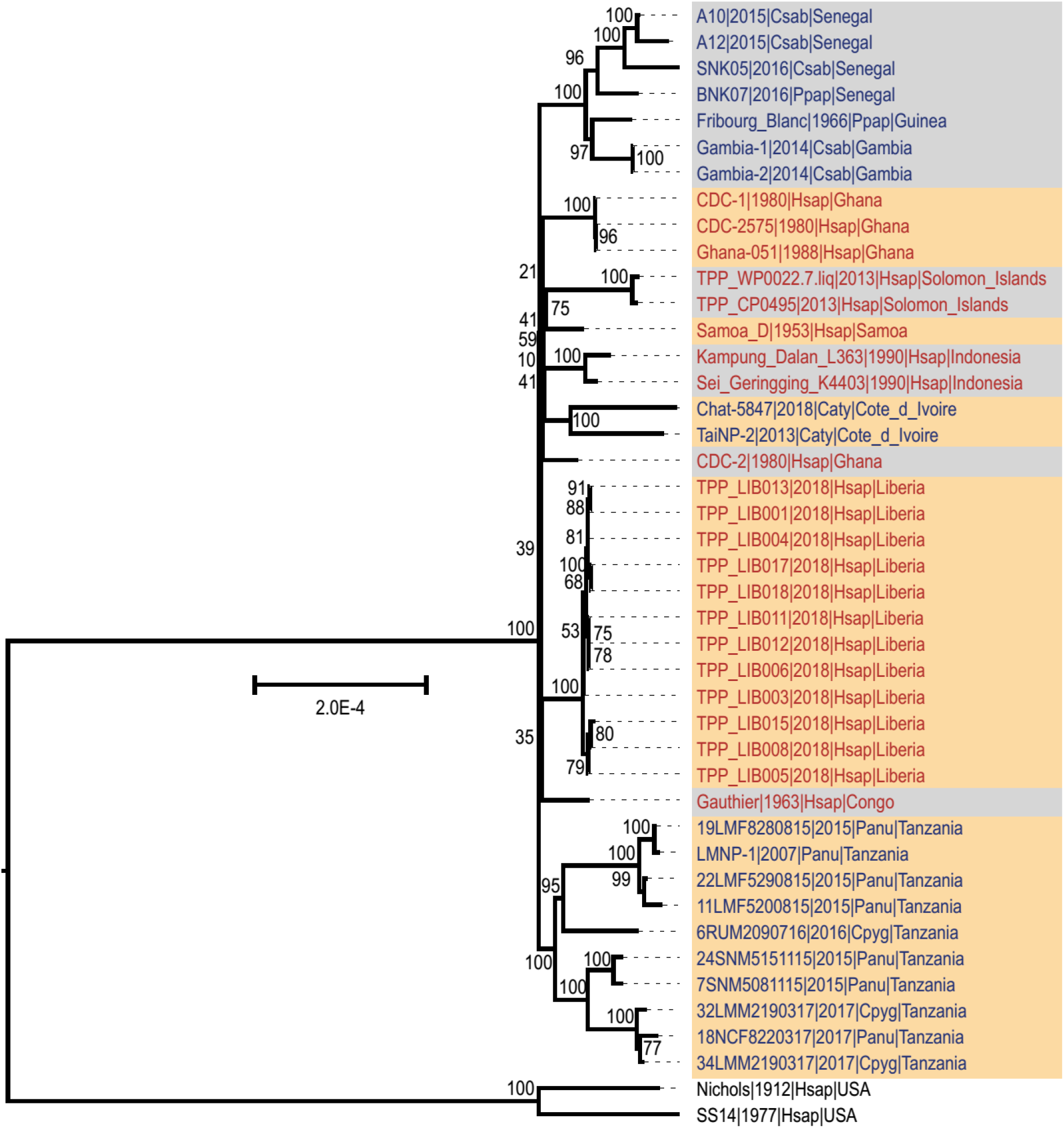
Maximum likelihood tree of *TPE*. The Nichols and SS14 samples are *TPA* samples and were used as an outgroup. *TPE* samples in blue and red represent isolates from **NHP** and humans, respectively. Gray and orange blocks delineate monophyletic clades with high branch support (bootstrap > 90) or isolated samples. The sample labels contain the lineage name, year of isolation, and country of isolation in this order.

The lack of evidence for recombination between *TPE* clades was also supported by analyses using the Gubbins and FastGear tools. In the case of Gubbins, the detected recombination signals were related to vertical transmission, involving only a parent node and its child node within the groups (https://github.com/sabrina-araujoo/Results/blob/main/core.full.aln.renamed.clean.recombination_predictions.gff). FastGear, which employs the BAPS algorithm to cluster samples before detecting recombination events, initially identified three distinct groups: two composed of *NHP* samples and one containing human isolates. However, in a subsequent analysis, FastGear merged all samples into a single lineage, possibly due to the low number of genomic variations among them.

Even without clear signs of recombination between human and non-human lineages, FastGear identified 11 recombinant regions in 9 samples, which may have originated from lineages not represented in this study (Table 2). Additionally, when the program was forced to maintain the initial segmentation of the three clusters defined by BAPS, a small recombinant region (between positions **149425 and 158147**) was detected in sample **18NCF8220317 (2017** | **Panu** | **Tanzania)**, suggesting a possible recombination event with a sample from the human cluster (Figure 4). Despite these findings, it is important to highlight that the low degree of genomic variability among the analyzed subspecies samples may represent a technical limitation, making it difficult to detect recombination events using the computational methodologies applied in this study.

**Table 2.** Recombinant regions inferred by FastGear originating from a lineage external to the.

| Start | End | log(BF) | StrainName |
| --- | --- | --- | --- |
| 230350 | 230589 | 0.1 | 24SNM5151115\|2015\|Panu\|Tanzania |
| 936384 | 936578 | 12.6 | 6RUM2090716\|2016\|Cpyg\|Tanzania |
| 230350 | 230589 | 0.1 | 7SNM5081115\|2015\|Panu\|Tanzania |
| 329918 | 330088 | 9.1 | BNK07\|2016\|Ppap\|Senegal |
| 372629 | 372754 | 63.7 | BNK07\|2016\|Ppap\|Senegal |
| 945382 | 948455 | 0.4 | TPP_WP0022.7.liq\|2013\|Hsap\|Solomon_Islands |
| 945382 | 948455 | 0.4 | TPP_CP0495\|2013\|Hsap\|Solomon_Islands |
| 936807 | 936894 | 7.5 | Kampung_Dalan_L363\|1990\|Hsap\|Indonesia |
| 523216 | 528892 | 11.5 | Chat-5847\|2018\|Caty\|Cote_d_Ivoire |
| 945074 | 946766 | 24.2 | Chat-5847\|2018\|Caty\|Cote_d_Ivoire |
| 940376 | 946766 | 6.5 | TaiNP-2\|2013\|Caty\|Cote_d_Ivoire |

**Figure 4.**
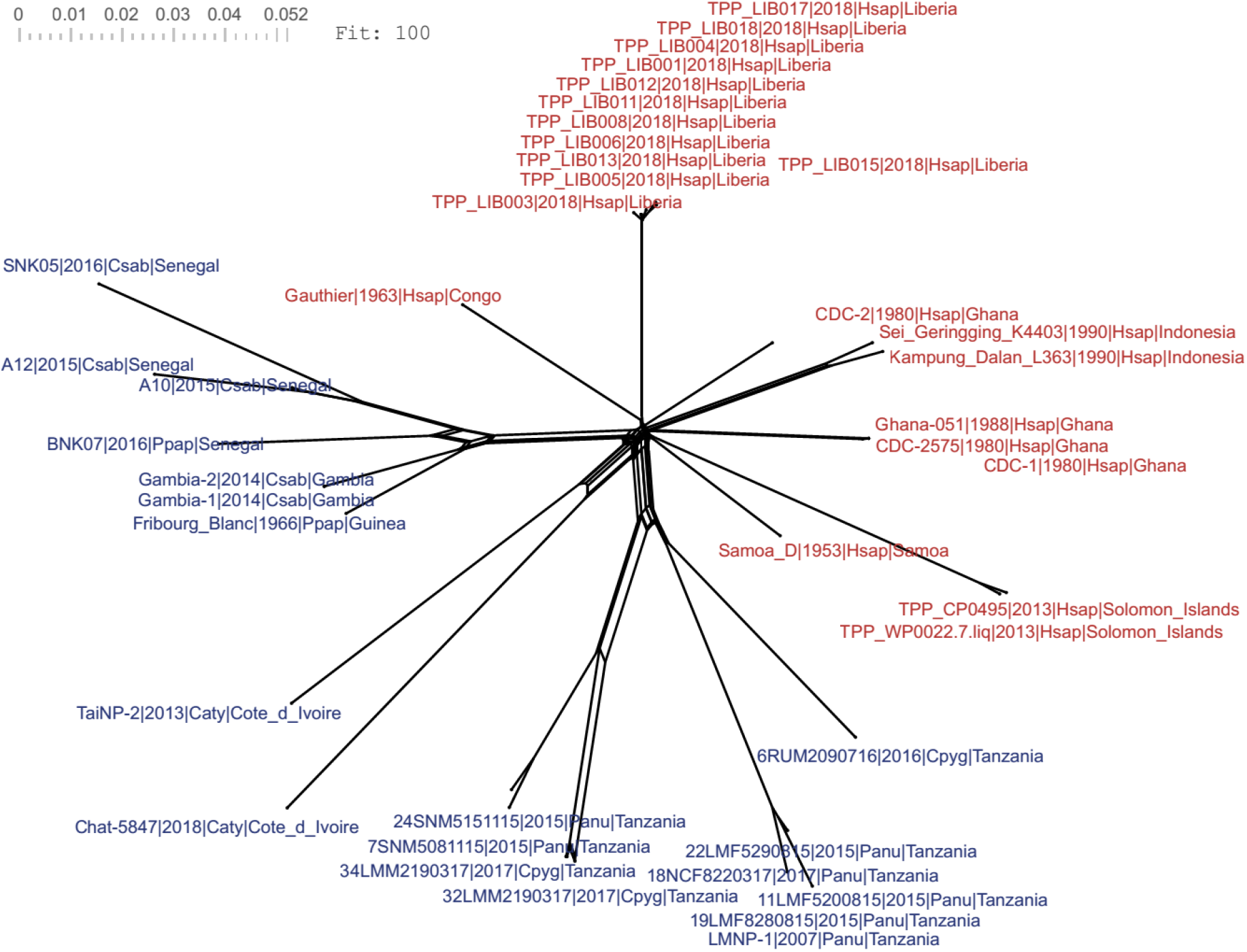
Reticulated tree generated by the SplitsTree program. Samples in blue and red indicate isolates from **NHP** and humans, respectively.

To complement the analysis, we performed a visual inspection of SNP alignments using a heatmap and hierarchical clustering methods to assess allele sharing between samples from different clades (Figure 5). The analysis demonstrates that the less frequent alleles at each site are mostly restricted to samples from one of the groups identified in the phylogenetic analysis.

**Figure 5.**
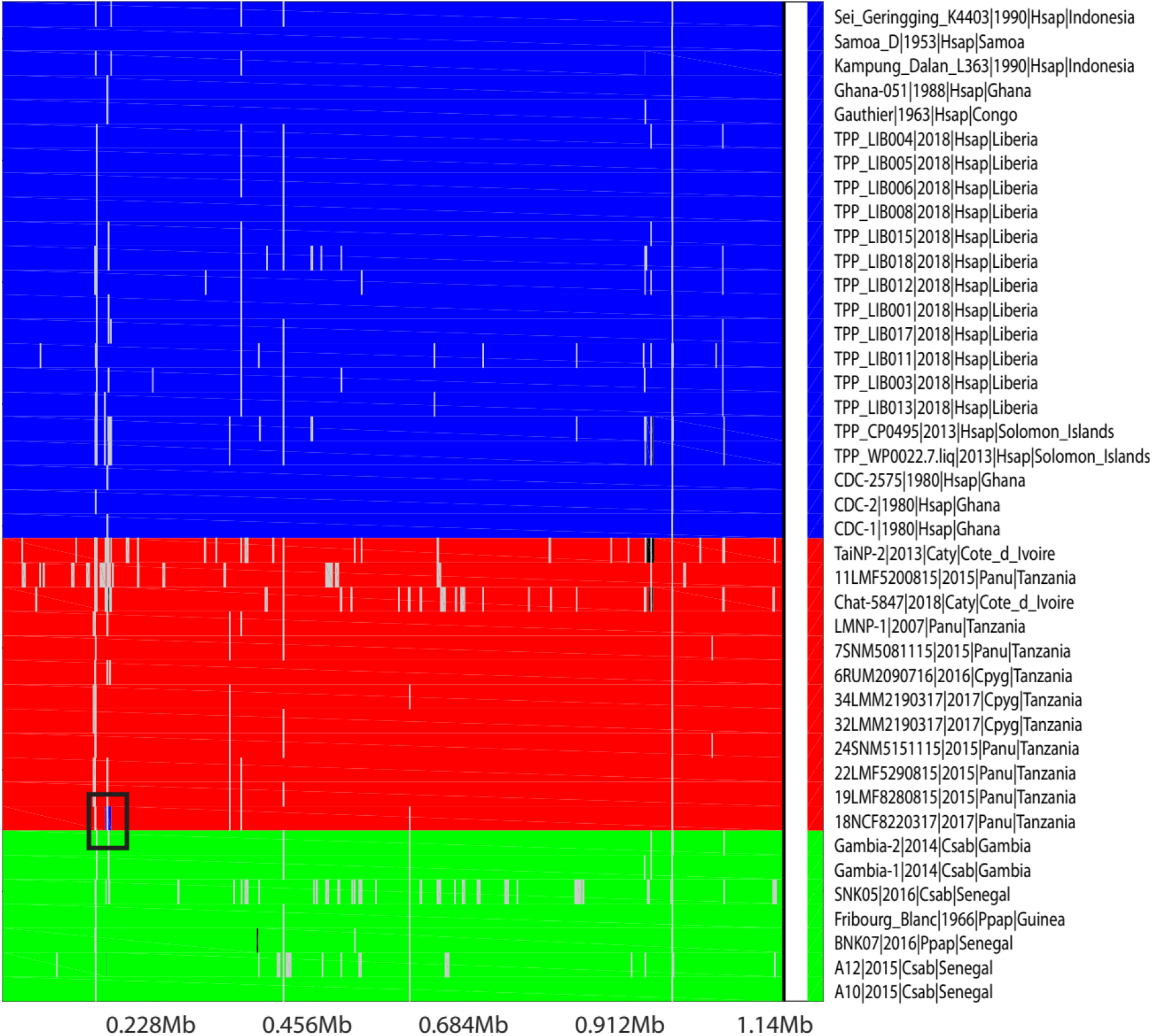
Inference of recombinant regions in *TPE* genomes performed by FastGear. Each row represents a sample, and the X-axis represents the genome length of the reference sample. The samples were grouped into three clusters by the BAPS program and are highlighted in blue, red, and green. The black box indicates a small region in sample **18NCF8220317** that may have received genetic material from a sample in the blue cluster. Gray and black regions represent, respectively, gap regions and recombinant regions originating from a lineage external to the samples in this study.

### 3.3. Origin and Evidence on the Host of the Common Ancestor of *T. pallidum* subsp. *pertenue*

The inference of the origin and ancestral host of *Treponema pallidum* subsp. *pertenue* was performed through the construction of a time-calibrated phylogenetic tree using the BEAST2 program. A preliminary analysis to verify whether the data followed a molecular clock structure, conducted with the TempEst program, yielded a low R^2^ statistic (0.137), indicating different mutation rate values across the branches of the tree. For this reason, we opted to use a relaxed molecular clock in the BEAST analysis, which allows each branch to have its own mutation rate. TempEst also indicated that the average mutation rate, measured by the slope, was 5.9E-7. The BEAST analysis included metadata regarding the year of isolation and the host of the samples (human or non-human primate), enabling a detailed reconstruction of the evolutionary trajectory of the subspecies. To ensure robustness in the results, two independent runs of the MCMC algorithm were conducted, both showing satisfactory convergence and ESS values above 200 for all inferred parameters, ensuring the reliability of the analysis.

The tree generated by BEAST2 (Figure 6A) exhibited a topology and clades with high support, consistent with the tree generated by IQ-TREE using the maximum likelihood method. However, an important distinction was observed: in the BEAST2 tree, the samples from the Solomon Islands and Samoa were grouped with high support, resulting in a total of nine well-defined clades. The internal nodes positioned at the base of the tree showed low support, again suggesting the rapid expansion of the subspecies during this evolutionary period. The estimated origin of the subspecies was dated to 1885 (1824–1932), indicating that its emergence and diversification occurred relatively recently on the evolutionary timescale.

**Figure 6:**
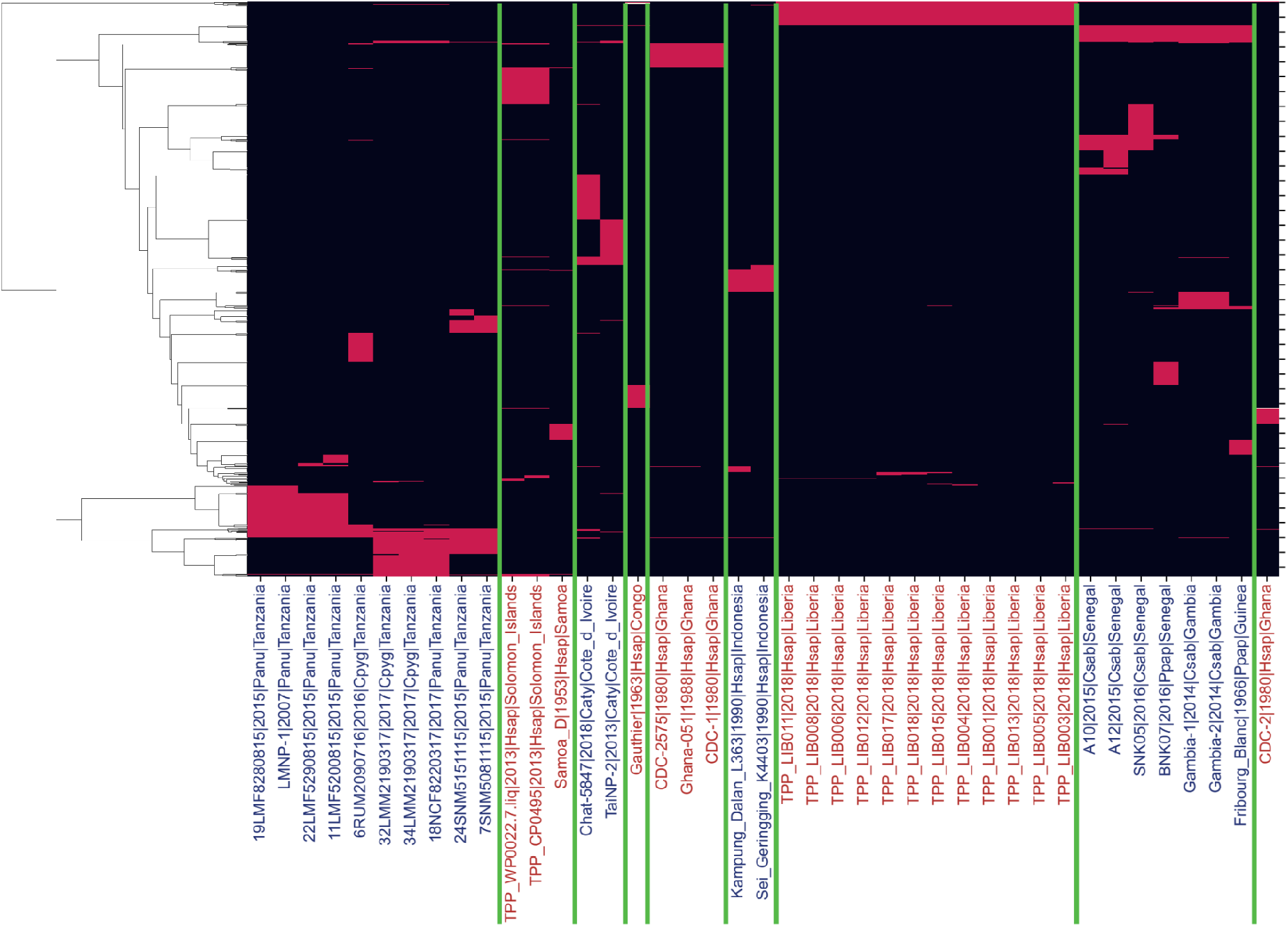
Heatmap highlighting polymorphic sites among the samples identified by the Snippy program. The rows and columns correspond to sites and samples, respectively. The less frequent alleles are shown in red, while the most frequent alleles are shown in black. The sites were clustered using hierarchical clustering with normalized mutual information as the distance metric and the Ward linkage method. The samples are arranged in the same order as presented in the ML tree (Figure 2), and vertical lines delineate the groups identified in the tree.

**Figure 7.**
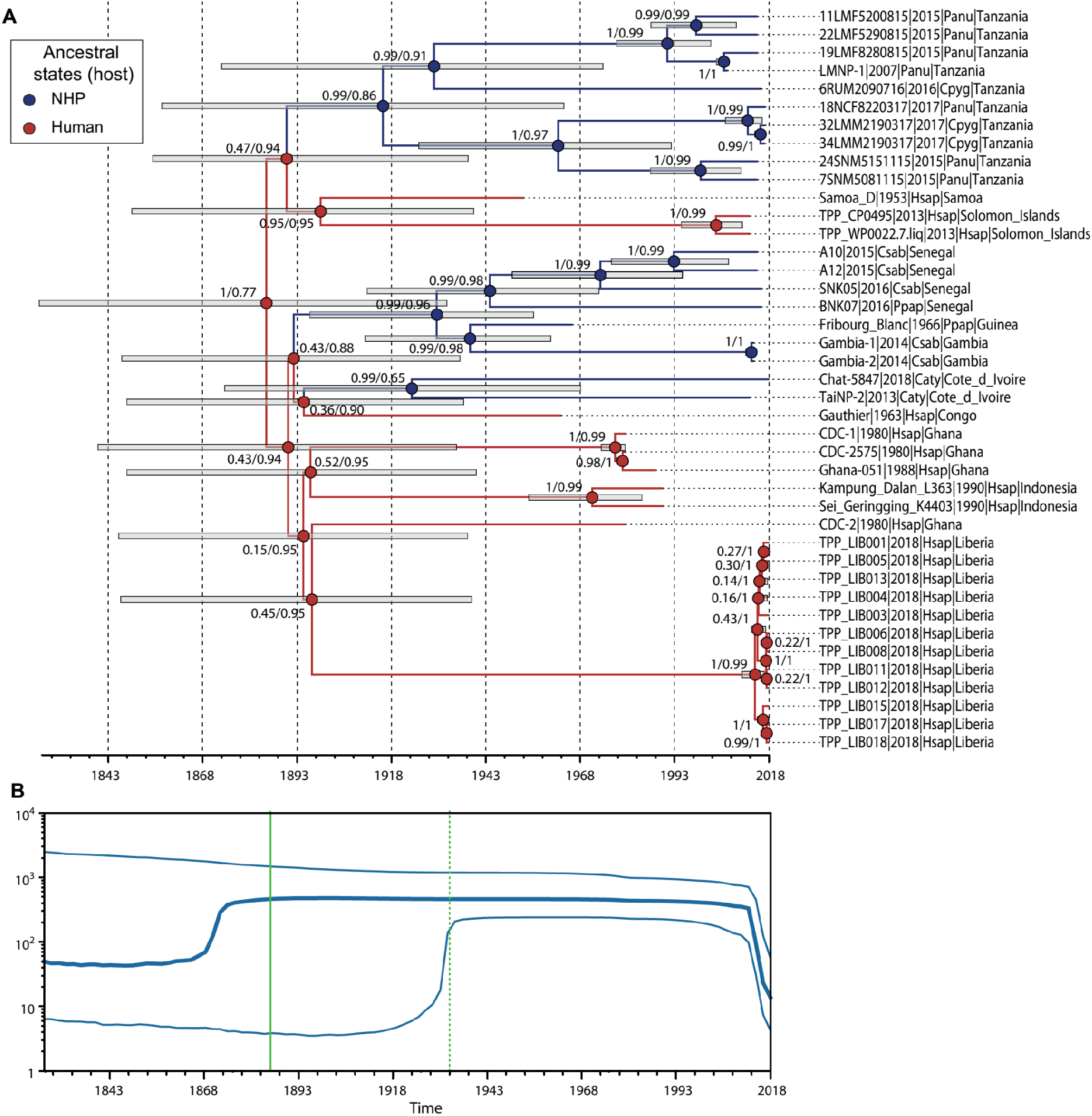
Bayesian inference on the evolution of TPE. MCC tree. The values on the internal nodes indicate, respectively, the posterior probability and the probability of the ancestral state. The colors of the branches and nodes correspond to the ancestral state with the highest probability (red: human; blue: NHP). The bars on the internal nodes represent the 95% HPD (Highest Posterior Density) of the node heights. B) Skyline plot describing the effective population size of TPE over time. The X-axis of the graph starts from the lower bound of the 95% HPD interval of the root node height (1824). The solid and dotted green vertical lines indicate, respectively, the mean height of the tree root (1885) and the upper bound of the 95% HPD interval of the root node height (1932) estimated in the Bayesian inference.

The ancestral host state analysis revealed that the most recent common ancestor of all analyzed samples had a high probability (77%) of having infected humans, reinforcing the hypothesis that the subspecies may have originated in human populations before potentially infecting non-human primates. Additionally, the Skyline Plot analysis (Figure 6B) demonstrated that the effective population size remained constant for a long period, around a value of 500. Subsequently, the subspecies underwent an abrupt reduction in its effective population size, possibly associated with yaws eradication campaigns, which significantly reduced the bacterium’s circulation among human hosts. These findings provide new insights into the evolution of *T. pallidum* subsp. *pertenue* and reinforce the importance of genomic monitoring to understand the transmission dynamics of this subspecies.

## 4. Discussion

In the present study, we aimed to deepen our understanding of the evolution of *Treponema pallidum* subsp. *pertenue* (TPE), with a focus on the origin and transmission dynamics of the disease, particularly in light of samples isolated from non-human primates (NHPs). The analyzed data were based on the study by JaneČková et al. (2023), whose previous investigations addressed various evolutionary aspects of this bacterium. Our approach complements these studies by exploring the possibility of transmission between humans and NHPs using additional computational tools, as well as inferring the probable host of the last common ancestor of TPE. We also highlight that some of the samples used by JaneČková et al. (2023) were filtered, retaining only those with high coverage. These analyses provide new perspectives on the ecological interactions and evolution of this pathogen, contributing to a broader understanding of its spread and adaptation among different hosts, as clinical observations have been increasing and gaining relevance since 1971 (Felsenfeld & Wolf, 1971; Fribourg-Blanc & Mollaret, 1969; Knauf et al., 2011; Chuma et al., 2018).

The origin of TPE in humans and non-human primates remains a topic of debate in the scientific literature, with evidence suggesting a common ancestral state among the different subspecies of *T. pallidum*. Phylogenomic studies indicate that the divergence between *T. pallidum* subspecies occurred from a common ancestor, possibly present in primate populations before being transmitted to humans (Harper et al., 2008; Korshunov et al., 2020). Whole-genome analyses reveal low genetic variability among *T. pallidum* strains, suggesting a relatively recent evolutionary event and strong selective pressure on the pathogen (Maděránková et al., 2019). Additionally, the presence of *T. pallidum* infections in non-human primates from different regions reinforces the hypothesis that *pertenue* may have emerged in wild environments before spreading to human populations, possibly through ecological interactions and close contact between host species (Knauf, Liu & Harper, 2013). However, the scarcity of genomic data, particularly from samples obtained directly from primates, hinders the precise reconstruction of the evolutionary history of this pathogen and prevents a detailed understanding of its transmission dynamics between humans and non-human primates. Thus, further studies with a larger number of samples and more robust phylogenomic approaches are essential to clarify the evolution and dispersion of *T. pallidum* subsp. *pertenue*.

The phylogenetic analysis conducted with IQ-TREE revealed clustering patterns within the species *Treponema pallidum* subsp. *pertenue* (TPE) similar to those reported by JaneČková et al. (2023), identifying ten groups, seven of which correspond to monophyletic clades with high statistical support. As observed in other studies, no clustering of samples isolated from humans and non-human primates was detected, and this clear separation between samples from different hosts suggests the existence of transmission barriers between them (Knauf et al., 2018).

Low resolution in the internal nodes at the base of the phylogenetic tree has also been observed in other studies (Knauf et al., 2018; JaneČková et al., 2023; Pla-Díaz et al., 2025). This low resolution may be attributed to a rapid expansion of the lineage in different hosts or the occurrence of genetic recombination events between clades, reinforcing the need for complementary studies to clarify this issue. The analyses conducted in this study using SplitsTree, Gubbins, and FastGear indicated that recombination events between sample groups are rare or nonexistent. The application of NeighborNet generated a reticulated tree with few reticulations between clades, supporting the hypothesis that vertical transmission predominates in the evolution of this subspecies. Furthermore, the Gubbins analysis confirmed that the detected recombination signals were restricted to vertical transmissions, with no evidence of significant horizontal gene flow between distinct hosts. FastGear initially identified three distinct groups, but upon refining the analysis, it merged all samples into a single lineage, possibly due to the low genomic diversity present in the subspecies. These results align with other studies that have examined recombination events among TPE samples (Knauf et al., 2018).

A more detailed analysis of the polymorphic sites found among TPE samples further demonstrated that the less frequent alleles were often restricted to one of the groups identified in this study, reinforcing the hypothesis that the low resolution at the base of the tree is due to the rapid expansion and isolation of ancestral TPE populations rather than recombination events.

Although recombination does not appear to have been a predominant factor in the evolution of TPE, 11 recombinant regions were detected in nine samples, suggesting the possible influence of lineages not represented in this study. Additionally, when maintaining the initial segmentation of clusters defined by BAPS in the recombination analysis using FastGear, a small recombinant region was identified in a sample isolated in Tanzania, raising the hypothesis of a recombination event involving a human clade—an issue previously suggested by Chuma et al. (2019). This result should not be overlooked, as the low genomic variability among the analyzed samples represents a technical limitation, making it difficult to accurately detect recombination events (Edmondson et al., 2021). Although the literature does not demonstrate the occurrence of recombination events among TPE samples, it is important to highlight that recombination events between *T. pallidum* subspecies have already been described (Pětrošová et al., 2012; Mikalová et al., 2017).

The time-calibrated phylogenetic reconstruction performed with BEAST2 provided additional insights into the evolutionary history of TPE. The low R^2^ statistic obtained from TempEst indicated heterogeneous mutation rates across the tree branches, justifying the choice of a relaxed molecular clock model (Pla-Díaz et al., 2025). The topology generated by BEAST2 was consistent with that inferred by IQ-TREE but revealed a newly well-supported clustering of samples from the Solomon Islands and Samoa, reducing the total number of well-defined groups to nine. The reduced support in the internal nodes reinforces the hypothesis of a rapid expansion of the subspecies during its initial period (Hackett et al., 1963). The analysis inferred that the origin of TPE dates to 1885 (1824–1932), indicating a recent emergence of TPE. Despite this inference, recent studies have detected the presence of *T. pallidum*, including TPE, in archaeological samples dating to periods earlier than those inferred in this study. A genome of this bacterium was detected in the remains of a Black Death victim in Lithuania, dated between the 15th and 17th centuries, suggesting that TPE was present in Europe during this period (Giffin et al., 2020). Although *Treponema pallidum* subsp. *pertenue* has traditionally been associated with tropical regions, recent studies suggest that variants of this bacterium may have circulated in Europe during historical periods, possibly due to trade interactions and population migrations (Harper et al., 2008; Giffin, 2020). Recent studies have also deepened our understanding of the presence of *Treponema* infections in pre-colonial Brazil, indicating that treponemal infections were already occurring in the region at least a thousand years before European contact. Based on bone samples found at an archaeological site in Santa Catarina, researchers identified DNA fragments of these bacteria and reconstructed their genome, indicating the presence of the bejel-causing bacterium (*Treponema pallidum* subsp. *endemicum*) in the region (Filippini, Pexo-Lanfranco & Edggers, 2019; Majander et al., 2024). This discovery challenges the traditional view that bejel was restricted to arid regions of the Old World, suggesting a broader and more complex geographical distribution of treponematoses. The discrepancy between the inferred dating of the origin from our phylogenetic analysis and the archaeological evidence is likely due to an overestimation of mutation rates, combined with the heterogeneous nature of these rates over evolutionary time. Phylogenetic analyses from recent studies, which include data from archaeological samples and other subspecies, indicate that the origin of *T. pallidum* predates the Common Era (over 5,000 years ago) and that the origin of TPE occurred more than 1,200 years ago (Pla-Díaz et al., 2025; Majander et al., 2024).

The inference of the ancestral host indicated a high probability (77%) that the most recent common ancestor of all analyzed samples infected humans, suggesting that the subspecies originated in human populations before spreading to non-human primates. This finding reinforces the theory previously proposed (Harper et al., 2008). Notably, this inference was conducted without including samples external to TPE, such as TPA and *T. pallidum* subsp. *endemicum*, both of which have humans as their host and could have biased the ancestral state inference toward humans. Among the literature reviewed, this represents the first study to perform this inference with statistical support. Additionally, the Skyline Plot analysis revealed a prolonged period of stability in the effective population size, followed by an abrupt decline, possibly associated with yaws eradication campaigns. These findings highlight the importance of genomic monitoring in understanding the transmission dynamics of TPE and reinforce the need for further investigations that integrate more comprehensive genomic analyses and functional experiments to elucidate interactions between different hosts. It is also important to note that TPE samples isolated from NHPs are currently restricted to African species. There have been no reports of *T. pallidum* infections in NHPs from other continents. It is of great interest to conduct studies that can detect TPE infections in NHP, particularly in New World primates. If evidence of *T. pallidum* infection is found in these individuals, it could significantly impact inferences regarding the evolution of *T. pallidum*.

